# Disentangling Rhythmic and Aperiodic Neural Activity: Investigating the Role of GABA Modulation in Sensorimotor Beta Oscillations

**DOI:** 10.1101/2025.02.17.638602

**Authors:** Jacopo Barone, Holly E. Rossiter, Suresh Muthukumaraswamy, Krish D. Singh

## Abstract

Beta oscillations (13–30 Hz) in the sensorimotor cortex are fundamental to motor control and are disrupted in motor disorders such as Parkinson’s disease and stroke. Animal studies and pharmacological research in humans have implicated GABAergic mechanisms in modulating these oscillations. Here, we explored the impact of GABAergic modulation on sensorimotor beta oscillations using magnetoencephalography (MEG) during a finger abduction task, with participants administered gaboxadol and zolpidem, two GABA-A positive allosteric modulators. We combined traditional oscillatory analyses with spectral parametrisation methods to separate periodic and aperiodic neural components. Time-frequency representations (TFRs) showed that gaboxadol induced a stronger modulation of beta dynamics than zolpidem, leading to deeper beta desynchronisation during movement and a more pronounced post-movement beta rebound. When accounting for aperiodic activity, gaboxadol showed modest effects limited to movement initiation, with increased beta power during movement onset but no significant impact during post-movement rebound. In contrast, zolpidem significantly enhanced beta power and altered aperiodic components, suggesting differential influences on GABAergic modulation and excitation-inhibition balance. These results underscore the necessity of distinguishing between periodic and aperiodic signals in spectral measures to improve the interpretation of neural oscillatory data.

## Introduction

Beta oscillations (∼13–30 Hz) are critical in motor control and are predominantly observed in the sensorimotor cortex and the basal ganglia, particularly in corticobasal ganglia-thalamic circuits (Liu et al., 2020; Sherman et al., 2016). Beta power is typically reduced during movement execution and increases during movement cancellation and error correction (Kilavik et al., 2013; Tan et al., 2016). While their full functional role remains under investigation, beta rhythms are thought to stabilise ongoing motor states and prevent premature actions by inhibiting new motor commands (Engel and Fries, 2010). Pathological alterations in beta rhythms are strongly associated with motor dysfunctions in conditions such as Parkinson’s disease and stroke, where excessive beta synchronisation correlates with bradykinesia, rigidity, and impaired motor recovery (Little and Brown, 2014; Thibaut et al., 2017). Additionally, in ageing, the decline in motor performance is often associated with reduced beta dynamics, suggesting a relationship between age-related changes in beta oscillations and motor decline (Rossiter et al., 2014).

Beta oscillations are closely tied to GABAergic interneuronal activity, which plays a central role in the generation of rhythmic patterns in the brain (Roopun et al., 2006; Whittington et al., 2000; Yamawaki et al., 2008). The relationship between beta oscillations and GABAergic signalling is supported by studies showing that pharmacological manipulation of GABA levels modulates beta activity in humans both during rest as well as during movement (Hall et al., 2010a; Muthukumaraswamy et al., 2013b; Nutt et al., 2015). Furthermore, evidence suggests that a reduction in cortical inhibitory tone, driven by GABAergic activity, is crucial for the induction of plasticity in the motor cortex—a process essential for motor learning (Bachtiar and Stagg, 2014). This strong link between beta rhythms and GABAergic inhibition highlights the intertwined role of these mechanisms in both normal and pathological motor control.

The dominant approach to analysing neural oscillations often focuses on changes in power within canonical frequency bands. In the frequency domain, oscillations appear as narrow-band peaks of power above the background aperiodic component (also known as 1/f). When changes in spectral power are observed, the implicit assumption is that frequency-specific modulations have occurred. However, a large portion of the signal recorded using LFP, EEG, and MEG is made up of irregular and arrhythmic activity, which is often overshadowed by the focus on oscillatory activity (He, 2014). Recent studies have highlighted the importance of this aperiodic activity, pointing to its significant alterations in various neuropsychiatric disorders (Bruining et al., 2020; Robertson et al., 2019) and the ageing brain (He et al., 2019; Voytek et al., 2015).

Recent pharmaco-EEG/MEG studies suggest that broadband spectral changes—often attributed to oscillatory shifts—could result from the underlying modulation of the 1/f component (Muthukumaraswamy et al., 2013b; Nutt et al., 2015). For example, pharmacological interventions targeting GABA-A receptors have been shown to modulate aperiodic activity, further complicating the interpretation of oscillatory power changes (Muthukumaraswamy et al., 2013a; Stock et al., 2020). These findings underscore the need to disentangle oscillatory (rhythmic) from non-oscillatory (aperiodic) components, as they likely originate from distinct neural mechanisms and serve different functional roles.

In this study, we investigated the link between the GABAergic system and sensorimotor beta oscillations, using a combination of MEG recordings and pharmacological interventions. We focused on the effects of gaboxadol and zolpidem during a finger abduction task. Although both drugs act as GABA-A positive allosteric modulators (PAMs), they engage distinct neural pathways. By separating the neural activity into periodic (rhythmic) and aperiodic (1/f) components, we aimed to demonstrate the importance of this distinction in understanding the specific effects of GABAergic modulation on various markers of neural activity.

## Results

### Behavioural Features

A series of paired *t*-tests were run to test the effect of the pharmacological intervention on behavioural performance. Reaction times were significantly slower after zolpidem intervention (*t*_(7)_ = −3.27, *p* = 0.014, *d* = −1.16, 95% CI [−2.18, −0.27]), suggesting a notable sedative impact. In contrast, no significant effect was found for gaboxadol (*t*_(8)_ = 0.262, *p* = 0.8, *d* = 0.09, 95% CI [−0.67, 0.98]) or placebo (*t*_(8)_ = −0.120, *p* = 0.9, *d* = −0.04, 95% CI [−0.7, 0.76]). Movement duration remained stable across all conditions: zolpidem (*t*_(7)_ = −1.69, *p* = 0.135, *d* = −0.6, 95% CI [−1.59, 0.38]), gaboxadol (*t*_(8)_ = 0.99, *p* = 0.349, *d* = 0.33, 95% CI [−0.53, 0.76]) and placebo (*t*_(8)_ = 0.5, *p* = 0.634, *d* = 0.17, 95% CI [−0.69, 0.89]). These findings indicate that zolpidem’s sedative effects significantly slowed reaction times, while gaboxadol and placebo had no measurable impact on behavioural responsiveness.

### Beta Power Modulation by Gaboxadol and Zolpidem

We investigated the effects of gaboxadol and zolpidem on average beta power using cluster-based permutation tests on the normalised TFRs, spanning the fulltime window from −1.5 to 1.5 seconds around movement offset. No significant clusters were detected between baseline sessions, indicating comparable pre-intervention conditions.

Time-frequency representations (TFRs) of oscillatory power are shown in Figure 1, where each plot represents the difference between post-intervention (POST60) and pre-intervention (PRE) spectra. Statistical analysis using cluster-permutation tests revealed that, compared to placebo, gaboxadol was associated with a significant cluster of decreased beta power over the left motor cortex (lM1) during the 500 ms window surrounding movement onset. The analysis also identified a significant power increase in the lM1 region during the 0.3 to 1 s post-movement window, indicating a stronger beta rebound after movement offset. For zolpidem versus placebo, cluster-permutation testing identified a significant reduction in beta power over lM1 within the 500 ms window around movement onset, while no significant clusters emerged in the post-movement period. When directly comparing gaboxadol and zolpidem conditions, statistical analysis revealed a significant cluster for lM1 in the window from −0.25 to 0.25 s around movement onset.

**Figure 1.**
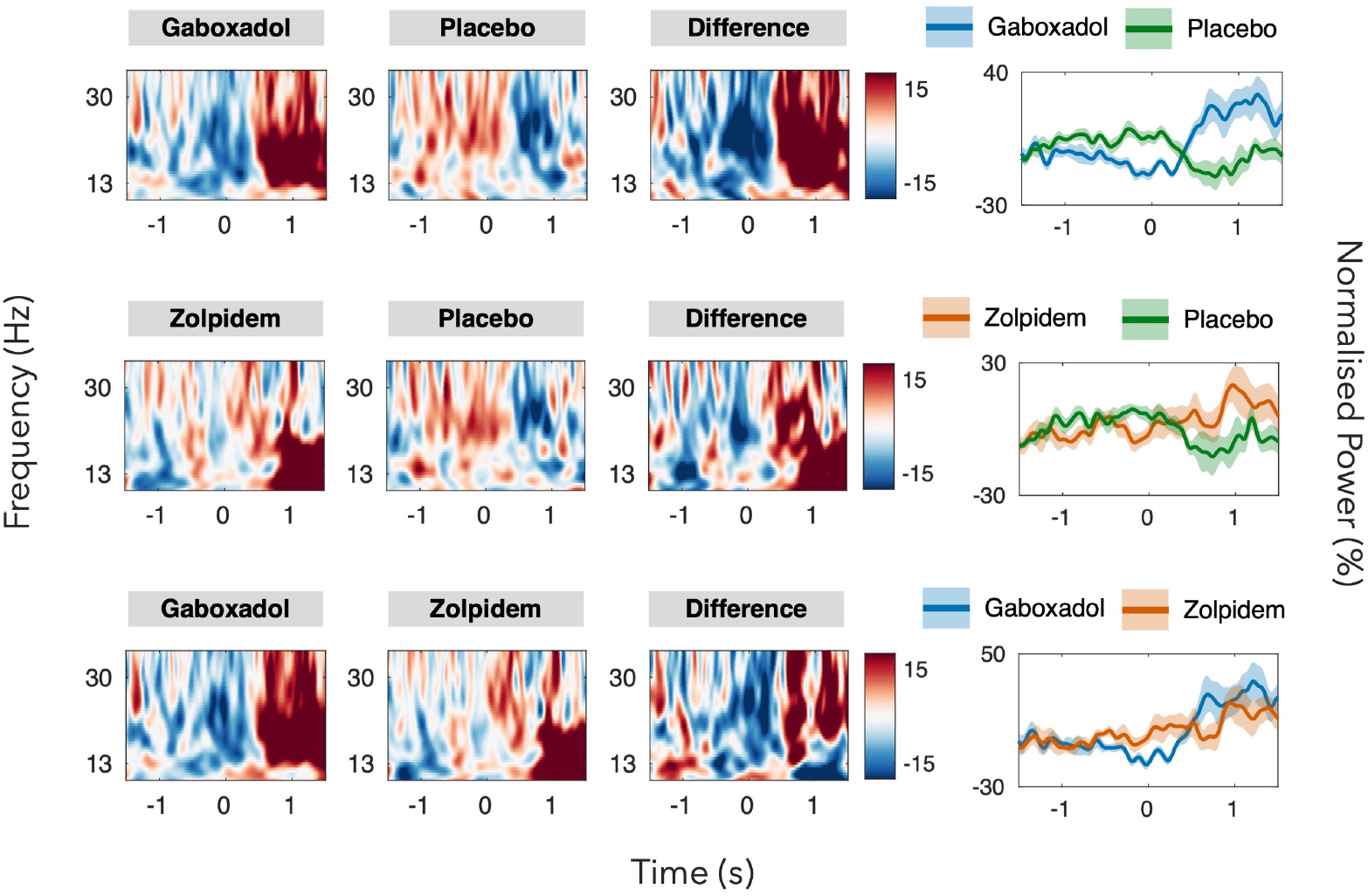
TFRs Comparison across conditions - POST60. The first two columns show time-frequency representations (TFRs) of normalised power differences (POST60 minus PRE) for the left primary motor cortex (lM1). The third column illustrates the contrast between conditions (difference of differences). Color scales represent changes in normalized power relative to baseline. The fourth column presents the temporal evolution of beta power (13–30 Hz), where solid lines show mean normalised power across participants and shaded areas indicate standard error (SE). Rows from top to bottom show: Gaboxadol vs. Placebo; Zolpidem vs. Placebo; and Gaboxadol vs. Zolpidem comparisons.

These results suggest that gaboxadol significantly modulates beta oscillations, enhancing beta dynamics both at movement onset and during the rebound period following movement. Gaboxadol appears to induce a more pronounced beta synchronisation at the start of the movement, followed by a stronger post-movement beta rebound. In contrast, zolpidem primarily affects beta synchronisation during movement, with less impact on the beta rebound.

### Effect of Pharmacological Intervention on Periodic and Aperiodic Features of Power Spectra

We further explored the results of averaged beta power by examining the dynamics of periodic and aperiodic aspects of power spectra. A two-way repeated measures ANOVA was conducted, with factors of pharmacological intervention (gaboxadol vs. zolpidem) and recording session (PRE vs. POST60). This analysis focused on spectra centred around a 250 ms window from −0.25 to 0 s around movement onset and from 0.75 to 1 s after movement termination, corresponding to the canonical time windows for beta desynchronisation and rebound respectively. We restricted our analysis to spectra from M1, which exhibited the largest and most consistent activation.

The ANOVA on aperiodic-adjusted beta power at movement onset revealed a significant main effect of intervention (*F*(1, 6) = 58.61, *p* < 0.001) and session (*F*(1, 6) = 12.78, *p* = 0.001) (Figure 2, *Top Row*). Post-hoc analysis indicated increased beta power at POST60 following gaboxadol (*t*(6) = 3.58, *p* = 0.021, *d* = 1.35, 95% CI [0.53, 3.62]) and zolpidem (*t*(6) = 3.01, *p* = 0.021, *d* = 1.17, 95% CI [0.33, 2.53]). Notably, zolpidem induced a larger increase in beta power compared to gaboxadol (*t*_(6)_ = 5.18, *p* = 0.004, *d* = 1.96, 95% CI [1.07, 3.62]). The ANOVA on aperiodic components revealed significant interactions for offset (*F*(1, 6) = 17.43, *p* = 0.006) and exponent (*F*(1, 6) = 23.76, *p* = 0.003) (Figure 2, *Bottom Row*). Post-hoc tests showed a significant decrease with zolpidem at POST60 for both aperiodic offset (*t*(6) = −5.90, *p* = 0.004, *d* = −2.23, 95% CI [−2.99, −1.49]) and exponent (*t*(6) = −5.75, *p* = 0.002, *d* = −2.17, 95% CI [−3.11, −1.17]). In contrast, gaboxadol did not yield significant modulations for offset (*t*(6) = 1.95, *p* = 0.147, *d* = 0.73, 95% CI [−0.22, 1.98]) and exponent (*t*(6) = 1.75, *p* = 0.131, *d* = 0.66, 95% CI [−0.28, 2.05]), although effects were in the opposite direction compared to zolpidem.

**Figure 2.**
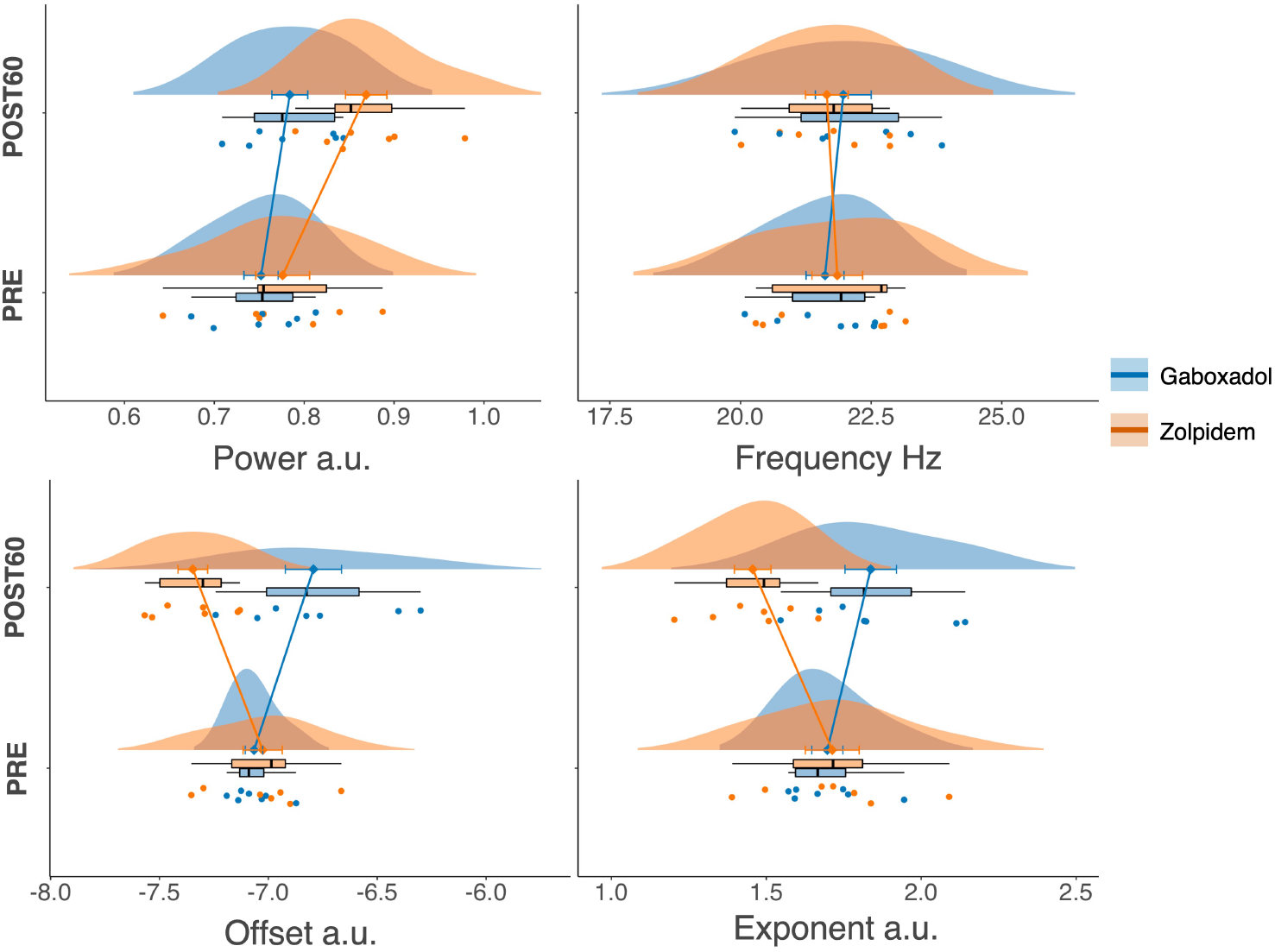
Comparison of gaboxadol and zolpidem interventions around movement onset in M1. Effects of pharmacological interventions on periodic (**beta power** – *upper left*, **beta frequency** – *upper right*) and aperiodic (**offset** – *lower left*, **exponent** – *lower right*) spectral features. Each raincloud plot displays individual participant scores (**coloured points**), boxplot, data distribution (**coloured curve**), and mean with standard error (**coloured diamond point and error bars**). Values are expressed in arbitrary units (a.u.). Observations are grouped by pharmacological intervention (light blue, Gaboxadol; light orange, Zolpidem) and session (PRE and POST60).

The ANOVA on beta power after movement termination indicated significant main effects of intervention (*F*(1, 6) = 13.14, *p* = 0.011) and session (*F*(1, 6) = 22.13, *p* =0.003) (Figure 3, *Top Row*). Paired *t*-tests revealed a substantial increase in beta power after zolpidem (*t*(6) = 5.71, *p* = 0.002, *d* = 2.16, 95% CI [1.57, 3.02]), while gaboxadol showed no significant effect (*t*(6) = 1.44, *p* = 0.2, *d* = 0.54, 95% CI [−0.27, 1.13]). The ANOVA on aperiodic offset revealed a significant main effect (*F*(1, 6) = 55.45, *p* < 0.001) and exponent (*F*(1, 6) = 29.95, *p* = 0.002), with significant post-hoc comparisons indicating an increase in aperiodic offset after gaboxadol (*t*(6) = 2.62, *p* = 0.04, *d* = 0.99, 95% CI [0.06, 3.6]) and a decrease after zolpidem (*t*(6) = −3.52, *p* = 0.04, *d* = −1.23, 95% CI [−1.92, −0.73]) (Figure 3, *Bottom Row*). The paired *t*-tests for the aperiodic exponent did not return significant findings after multiple comparisons for either gaboxadol (*t*(6) = 1.71, *p* = 0.14, *d* = 0.65, 95% CI [−0.21, 2.52]) or zolpidem (*t*(6) = −2.86, *p* = 0.06, *d* = −1.08, 95% CI [−1.92, −0.67]).

**Figure 3.**
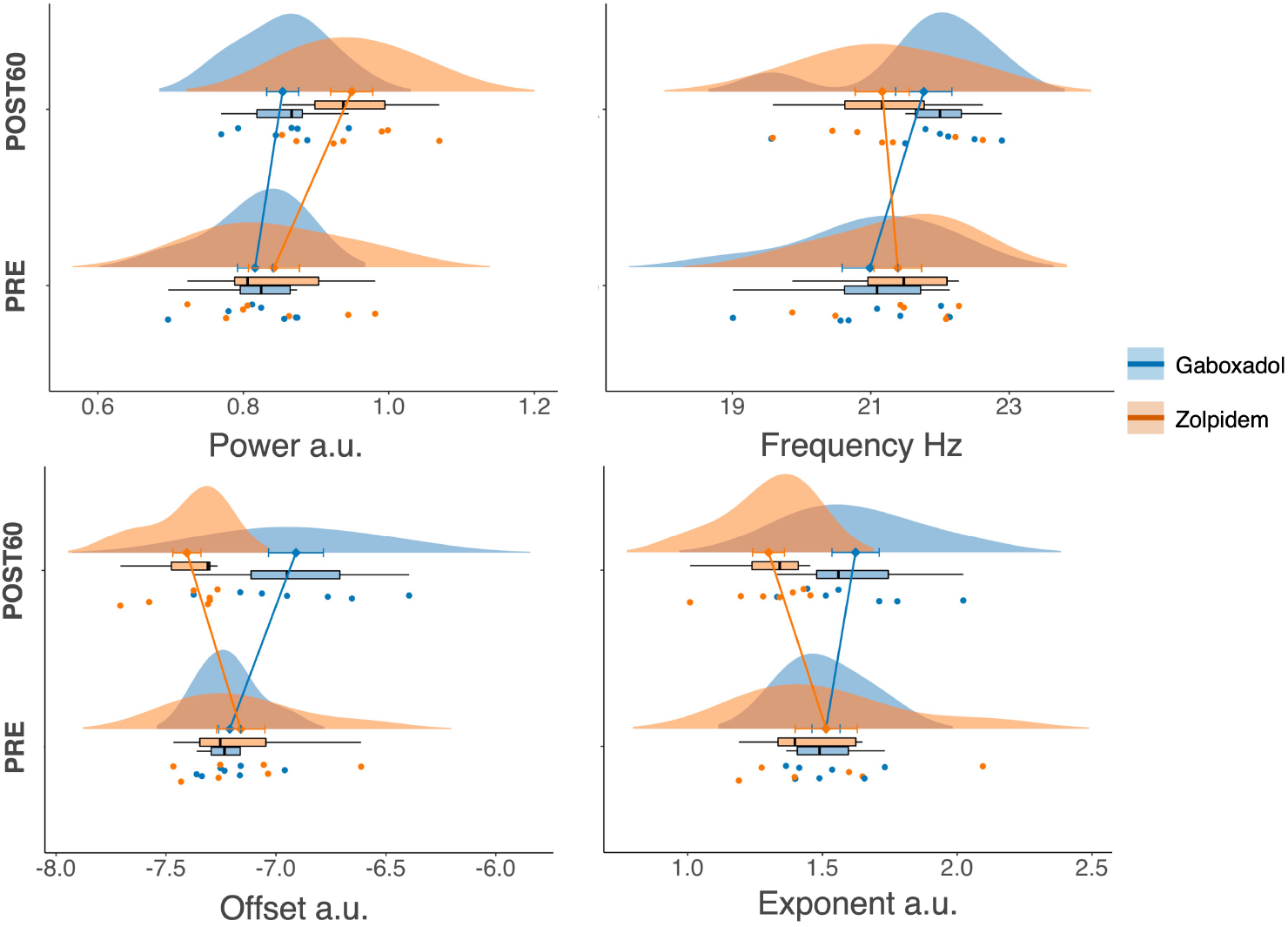
Comparison of gaboxadol and zolpidem interventions around movement offset in M1. Effects of pharmacological interventions on periodic (**beta power** – *upper left*, **beta frequency** – *upper right*) and aperiodic (**offset** – *lower left*, **exponent** – *lower right*) spectral features. Each raincloud plot displays individual participant scores (**coloured points**), boxplot, data distribution (**coloured curve**), and mean with standard error (**coloured diamond point and error bars**). Observations are grouped by pharmacological intervention (light blue, Gaboxadol; light orange, Zolpidem) and session (PRE and POST60).

In summary, the pharmacological interventions had distinct effects on the brain’s beta oscillations and underlying neural activity patterns. Gaboxadol, while enhancing beta power during movement initiation, had a weaker influence on the rebound following movement. Zolpidem, on the other hand, led to a stronger beta power increase and significant changes in the non-oscillatory, or aperiodic, components of the neural signal. These results indicate that these drugs modulate both the oscillatory and aperiodic features of neural activity in different ways, with zolpidem having a more pronounced effect on the overall neural signal structure.

## Discussion

In this study, we examined the effects of two GABAergic modulators, gaboxadol and zolpidem, on sensorimotor beta oscillations. By using spectral parametrisation, we were able to differentiate the complex effects of these pharmacological interventions on rhythmic neural activity, providing a more nuanced understanding of the underlying physiology.

We observed selective effects on periodic spectral components following the interventions. Zolpidem consistently increased aperiodic-adjusted beta power, whereas gaboxadol had only a marginal impact. Interestingly, these findings diverge somewhat from the averaged beta power results. Specifically, time-frequency representations (TFRs) showed that gaboxadol induced a stronger modulation of beta dynamics than zolpidem, leading to deeper beta desynchronisation during movement and a more pronounced post-movement beta rebound (PMBR). This discrepancy may arise because rhythmic neural activity is intertwined with aperiodic components, as has been previously reported in the literature (Donoghue et al., 2020). Aperiodic activity is highly dynamic and has been associated with contexts typically studied for oscillatory phenomena, such as cognitive tasks or disease states (Bruining et al., 2020; Lendner et al., 2020; Molina et al., 2020). Traditional power decomposition methods, which focus on band-pass filtered signals, may misrepresent oscillatory components by conflating them with aperiodic signals, potentially leading to erroneous interpretations of the physiological processes at play (Donoghue et al., 2020, 2022; Gerster et al., 2022).

Our study revealed bi-directional aperiodic modulations following pharmacological intervention. Zolpidem reduced both the aperiodic offset and exponent, whereas gaboxadol increased these parameters. The aperiodic offset has been linked to broadband amplitude dynamics, reflecting cortical activity at the population level and is associated with increased neuronal spiking (Miller et al., 2014; Manning et al., 2009). Meanwhile, the aperiodic exponent is considered a proxy for the excitation-inhibition E: I balance, with higher values suggesting increased inhibitory (GABAergic) activity and lower values indicating a shift towards excitation (Gao et al., 2017; Miller et al., 2009). These differential effects on aperiodic components align with previous work showing that distinct GABAergic mechanisms can produce specific and measurable changes in the aperiodic exponent, supporting its utility as a marker of E:I balance (Muthukumaraswamy et al., 2013a; Muthukumaraswamy and Liley, 2018; Robertson et al., 2019; Stock et al., 2020).

Previous pharmaco-EEG/MEG studies investigating the GABAergic system in humans have consistently reported an increase in beta power after GABA-A agonist or GABA reuptake blocker administration, both at rest and during movement (Hall et al., 2010a, 2011; Jensen et al., 2005; Muthukumaraswamy et al., 2013b). Studies using gaboxadol and zolpidem have similarly demonstrated effects on beta oscillations, though distinct spectral profiles were observed for each drug (Nutt et al., 2015). Our findings partially align with these reports. Gaboxadol’s effects on beta oscillations were modest, appearing primarily around movement onset, with no significant influence on post-movement rebound. This pattern resembles findings from Hall et al. 2011, where diazepam enhanced beta desynchronisation without modulating PMBR. In contrast, Muthukumaraswamy et al. 2013b observed modulation of both desynchronisation and PMBR with tiagabine, a GABA reuptake blocker that acts on both GABA-A and GABAB receptors. Our findings suggest that gaboxadol, which like diazepam targets GABA-A receptors, may have a more selective effect on beta desynchronisation, supporting the hypothesis that desynchronisation and rebound are distinct phenomena with different underlying mechanisms (Gaetz et al., 2011; Jurkiewicz et al., 2006; Muthukumaraswamy et al., 2013b).

Zolpidem, on the other hand, produced a marked increase in beta power. This is consistent with findings from rodent models where zolpidem increased beta oscillations by enhancing phasic interneuron activity (Yamawaki et al., 2008), and with a human MEG study that reported zolpidem-mediated increases in beta power (Nutt et al., 2015). However, studies involving patients with atypical beta oscillations, such as those with Parkinson’s disease or stroke, suggest that subsedative doses of zolpidem can reduce beta power and improve motor and cognitive functions, likely due to its specific action on GABAergic projections in the basal ganglia (Hall et al., 2010b; Prokic et al., 2019). This contrasts with the increase in beta power observed in our study, which may be related to the sedative effects of the higher dose used.

We propose that changes in aperiodic activity can partially explain these complex GABAergic effects on beta oscillations. Standard signal decomposition methods, such as wavelet and short-time Fourier transforms, do not explicitly account for the aperiodic components, potentially confounding the detection and interpretation of genuine oscillatory activity (Donoghue et al., 2020). Gaboxadol’s increased aperiodic offset and exponent suggest a rise in local neuronal activity and heightened inhibition, which may contribute to increased low-frequency power in the spectrum, even in the absence of enhanced rhythmic activity. In contrast, zolpidem’s reduction of both aperiodic components likely corresponds to decreased low-frequency and increased high-frequency power, resulting in a flattened spectrum. This spectral flattening aligns with previous reports of zolpidem enhancing beta and low-gamma power while reducing alpha power (Nutt et al., 2015).

It is crucial to interpret these findings cautiously, particularly given the challenges posed by the sedative effects of the pharmacological interventions, which likely influenced behavioural performance. In our study, slower reaction times were observed after zolpidem administration. To separate the effects of behaviour on our results, we considered comparing conditions using a subset of trials with similar reaction times. However, this approach was not feasible due to the limited number of trials and the consistent performance issues caused by the interventions. Future studies could address this limitation by controlling for behavioural variability or increasing the number of trials to ensure robust statistical comparisons. While our study focused on spectral parametrisation, it would greatly benefit from complementary analyses, such as investigating beta bursts. Recent work, such as the study by Seymour et al. 2022, has shown that combining burst analysis with spectral parametrisation offers a more comprehensive view of oscillatory dynamics. Incorporating aperiodic activity into burst analyses could improve sensitivity to detect transient oscillatory bursts and provide deeper insights into the interaction between GABAergic modulation and beta rhythms.

In summary, we emphasize the importance of distinguishing between periodic and aperiodic components in electrophysiological recordings to avoid misinterpretation of neural oscillations. Advances in methods for decomposing signals and separating these components provide powerful tools for understanding the true nature of neural rhythms and the physiological processes they reflect (Donoghue et al., 2020, 2022; Gerster et al., 2022; Kosciessa et al., 2020; Wen and Liu, 2016; Wilson et al., 2022).

## Methods

### Design and participants

Twelve healthy male participants (mean age 27.7, range 21-35) took part in a randomised, single-blind, placebo-controlled study comparing single doses of zolpidem and gaboxadol. The dataset was collected at Cardiff University Brain Research Imaging Center by Prof. Suresh Muthukumaraswamy. These data were collected during the same recording sessions as Nutt et al. 2015, but the specific analyses presented in this paper have not been previously published.

Doses of 15 mg gaboxadol and 10 mg zolpidem were chosen. This was well-tolerated and showed comparable sedative properties (Hajak et al., 2009). The elimination half-lives of these two drugs were similar at approximately 1.5-2.0 h in fasting subjects. All participants were medically screened and excluded for significant medical, psychiatric or neurological conditions and current recreational or prescription drug use. For each session, participants were scanned on separate days, after at least a seven-day washout period, at approximately the same time of day. On each day, an initial baseline MEG recording was obtained (PRE). Participants then orally ingested a capsule containing either a placebo, 15 mg of gaboxadol or 10 mg of zolpidem. Participants were blinded to the contents of the capsules and the placebo/control session order was counterbalanced across both experiments. MEG recordings were obtained at a 60-minute time point (POST60). Zolpidem was sourced from an NHS hospital pharmacy, and gaboxadol was donated by Lundbeck as part of the ECNP Medicines Chest Initiative (Nutt et al., 2014).

For each MEG recording, the participants performed 100 trials of a cued finger movement task, comparable to that described in Muthukumaraswamy, 2010. The task was chosen because of its simplicity and the ability to modulate beta power effectively. The participants were required to perform ballistic abductions of the right-hand index finger to an auditory tone pip played through insert headphones (4.5 s ISI). The participants’ index fingers were lightly attached to a piece of plastic to measure finger displacement. To maintain a constant motor performance throughout the experiment the following feedback procedure was implemented. After the auditory pip (1.5 s), the participants received on-screen feedback with a “virtual ruler”, indicating how far they had moved relative to a target movement criterion (10 mm). This feedback stayed on the screen for 1 second and then was replaced with a fixation cross. The participants quickly learned to move consistently on each trial and had training trials at the beginning of each day.

### MEG acquisition and preprocessing

MEG signals were recorded using a CTF whole-head system with 275 axial gradiometer channels configured in a synthetic third-order gradiometer mode. The signals were recorded at a sampling rate of 1200 Hz. Simultaneous EMG recordings were made from the participants’ right first dorsal interosseus (FDI) and digitised with the MEG data. The participants’ fingers were lightly attached to a small piece of plastic, attached to an optical displacement system. This device gave a one-dimensional measure of displacement (in the direction of index-finger abduction), which was also continuously sampled with the MEG. Fiduciary coils were placed at fixed distances from three anatomical landmarks (nasion, left, and right pre-auricular) and the positions of the coils were monitored continuously throughout the session. Each participant had a 1 mm isotropic FSPGR MRI scan available for source localisation analysis. To achieve MRI/MEG co-registration, the fiduciary markers were placed at fixed distances from anatomical landmarks identifiable in the participants’ anatomical MRIs (tragus, eye centre). The MEG data were acquired continuously and epoched offline. All analyses were performed in MATLAB (MathWorks Inc, Natick, MA), mainly using the FieldTrip toolbox (Oostenveld et al., 2011) and custom scripts. MEG signals were first high-pass and low-pass filtered at 0.5 Hz and 150 Hz respectively. Spectral interpolation was used to remove powerline contamination and harmonics (Leske and Dalal, 2019). Data trials including large muscle artefacts were identified via a semi-automatic procedure. Trials were band-pass filtered between 110-140 Hz, z-transformed and compared against a threshold. Trials with values above the cut-off were visually inspected before exclusion. Eye movements and cardiac artefacts were projected out of the data using independent component analysis (Makeig et al., 1995). Finally, MEG signals were down-sampled to 300 Hz. From the continuous MEG recordings, EMG onsets were marked using an automated algorithm that marked increases in the rectified EMG signal by 1.5 SD above the noise floor, subject to the constraint that they occurred within 750 ms of the tone pip. Data were epoched from −1.5 s to 1.5 s around the start (EMG onset) and the end (EMG offset) of the movement.

Gaboxadol recordings were not available for two participants, while three participants were excluded for zolpidem. Another participant was excluded due to a high number of faulty trials in all recordings, caused by a combination of muscle artefacts and poor performance. The number of participants available after preprocessing was 9 for gaboxadol, 8 for zolpidem and 11 for placebo. Trials were adjusted at the end of each recording so that each participant had an equal number of trials between sessions (PRE, POST60). Furthermore, pharmacological interventions were contrasted in pairs (gaboxadol vs placebo; zolpidem vs placebo; gaboxadol vs zolpidem). The number of participants tested was balanced to allow within-subject comparisons.

### Source imaging

For source localisation, each participant’s anatomical MRI was divided into irregular grid by warping the individual MRI to the MNI template brain and then applying the inverse transformation matrix to the regular MNI template grid (4mm isotropic voxel resolution), allowing source estimates at brain locations directly comparable across participants. For each grid location inside the brain, the forward model (i.e. the lead field) was calculated for a single dipole orientation by singular value decomposition, using a single-shell volume conduction model (Nolte, 2003). Since all grid locations of each subject were aligned to the same anatomical brain compartments of the template, corresponding brain locations could be statistically compared over all subjects. Source power at each location was estimated using an LCMV (linearly constrained minimum variance) beamformer (Van Veen et al., 1997). Beamformer analysis uses an adaptive spatial filter to estimate the power at every specific (grid) location of the brain. Virtual time courses were reconstructed for a set of cortical ROIs. For each ROI, the virtual time courses with the largest SD across time were selected as the target virtual channel for the ROI.

A data-driven pipeline was employed to extract relevant ROIs. Sources were reconstructed in the beta band for the placebo intervention, with a frequency domain beamformer source analysis performed by using the dynamic imaging of coherent sources algorithm (DICS) (Gross et al., 2001). The spatial filter was constructed from the individual lead fields and the cross-spectral density (CSD) matrix for each subject. CSD matrices were computed for the task period ranging from 0 to 500 ms after the auditory tone and a baseline period of the same length, with an offset of −500 ms relative to the auditory tone. CSD matrices were computed in the beta band for 25 Hz (± 10 Hz) where spectral smoothing is indicated in brackets. CSD matrix calculation was performed with the multitaper method (Percival and Walden, 1993) using four Slepian tapers (Slepian, 1978). An activation-versus-baseline *t*-statistic was calculated at a single participant level by using an analytic dependent-samples within-trial *t*-test. The source *t*-values obtained were grouped in ROIs according to the AAL atlas in FieldTrip (Tzourio-Mazoyer et al., 2002). Then *t*-values were thresholded at alpha = 0.05 and the proportion of significant sources for each ROI was computed. This process was repeated for both the PRE (no drug) and the POST60 sessions. The top 10% ROIs with the highest proportion of significant sources between the two sessions were selected for extracting virtual time courses. The ROIs list according to the AAL atlas was composed of the Precentral cortex (M1), Postcentral cortex (S1), Paracentral Lobule (PL), Mid Cingulum (mC) and Supplementary motor area (SMA).

### TFRs on virtual channel time courses

Preprocessed MEG signals were decomposed into their time-frequency representations (TFRs) in the 10-35 Hz range using a Hanning taper with a sliding time window of 7 cycles. MEG power change was subsequently normalised as the percentage change relative to the overall average by dividing the power at each frequency and each time point by the average power of that frequency across the whole experimental session (Tan et al., 2016; Torrecillos et al., 2015). Values *>*0 indicated power higher than the overall average power of that frequency and vice versa.

### Statistical analysis

The pre-intervention baseline spectra (PRE) were subtracted from each post-intervention spectra (POST60) and then differences between interventions were tested at two different latencies: from −0.25 to 0.25 s around movement onset and from 0.3 to 1 s after movement offset. For this purpose, we used a dependent-samples permutation *t*-test and a cluster-based correction method (Maris and Oostenveld, 2007) to account for multiple comparisons across frequencies (Monte Carlo estimate). Samples whose *t*-values exceeded a threshold of cluster *α* = 0.05 were considered as candidate members of clusters of adjacent samples. The sum of *t*-values within every cluster was calculated as test statistics. These cluster sizes were then tested (two-sided) against the distribution of cluster sizes obtained for 10000 repetitions.

Repeated-measures ANOVAs were used to investigate the effects of pharmacological interventions (gaboxadol vs zolpidem) and experimental sessions (PRE vs POST60). Mauchly’s test of sphericity was used to test the homogeneity of variance. Where Mauchly’s test of sphericity was significant (p <0.05) in repeated-measures ANOVAs, Greenhouse-Geisser corrections were applied. Two-tailed paired-sample *t* tests were calculated using FDR correction for multiple comparisons. Effect sizes were calculated using Cohen’s d, calculated as the difference between the two means, divided by the standard deviation of the difference. 95% confidence intervals (95% CI) were calculated using accelerated bias-corrected percentile limits (number of bootstrap samples = 10000).

### Power spectra parametrisation

The Spectral Parameterisation Resolved in Time algorithm (SPRiNT) (Wilson et al., 2022) is designed to identify and model spectral features of neural activity across time. First, the algorithm performs a short-time Fourier transform (STFT) on 0.5 s sliding time windows using MATLAB’s FFT. Time windows are then averaged locally in time (3 windows; 50% overlap) to generate local-mean power spectra. Power spectra are then parametrised implementing the same algorithm used in the FOOOF toolbox (Donoghue et al., 2020). The toolbox conceptualises the power spectra as a combination of an aperiodic component with overlying periodic components (oscillations). These putative oscillatory components are characterised as frequency regions of power over and above the aperiodic component. The aperiodic component is fit as a function across the entire fitted range of the spectrum, and each oscillatory peak is individually modelled with a Gaussian. The final outputs of the algorithm are the parameters defining the best fit for the aperiodic component and the Gaussians. These are described by the exponent and offset, and periodic peaks, are described by the centre frequency, power, and bandwidth of identified peaks/Gaussians. The FOOOF algorithm was called with the following settings: frequency range 3-40 Hz; peak width limits 1.5-6; max peaks 4; min peak height 0.2; aperiodic mode fixed; peak threshold 2.

